# The TNF^ΔARE^ mouse as a model of intestinal fibrosis

**DOI:** 10.1101/2023.01.13.523973

**Authors:** Calen A Steiner, Samuel D Koch, Tamara Evanoff, Nichole Welch, Rachael Kostelecky, Rosemary Callahan, Emily M Murphy, Caroline H T Hall, Sizhao Lu, Mary CM Weiser-Evans, Ian M Cartwright, Sean P Colgan

## Abstract

**Background & Aims:** Crohn’s disease (CD) is a highly morbid chronic inflammatory disease. The majority of CD patients also develop fibrostenosing complications. Despite this, there are no medical therapies for intestinal fibrosis. This is in part due to lack of high-fidelity biomimetic models to enhance understanding and drug development. There is a need to develop *in vivo* models of inflammatory bowel disease-related intestinal fibrosis. We sought to determine if the TNF^ΔARE^ mouse, a model of ileal inflammation, may also develop intestinal fibrosis.

**Methods:** Several clinically relevant outcomes were studied including features of structural fibrosis, histological fibrosis, and gene expression. These include the use of a luminal casting technique we developed, traditional histological outcomes, use of second harmonic imaging, and quantitative PCR. These features were studied in aged TNF^ΔARE^ mice as well as in cohorts of numerous ages.

**Results:** At ages of 24+ weeks, TNF^ΔARE^ mice develop structural, histological, and genetic changes of ileal fibrosis. Genetic expression profiles have changes as early as six weeks, followed by histological changes occurring as early as 14-15 weeks, and overt structural fibrosis delayed until after 24 weeks.

**Discussion:** The TNF^ΔARE^ mouse is a viable and highly tractable model of intestinal fibrosis. This model and the techniques employed can be leveraged for both mechanistic studies and therapeutic development for the treatment of intestinal fibrosis.

## INTRODUCTION

Inflammatory bowel disease (IBD) is a family of disorders characterized by chronic, relapsing inflammation of the gastrointestinal tract^1^. The prevalence of IBD varies worldwide and corresponds roughly with the degree of industrialization; as a result, developed countries show the highest rates of IBD, but with accelerating prevalence rates in the developing world due to burgeoning industrialization^2^. IBD is currently sub-divided into two major classes, Crohn’s Disease (CD) and ulcerative colitis (UC). There are over 780,000 patients with Crohn’s disease (CD) in the US, and most will eventually develop fibrostenotic complications of the intestine.^3^ Billions of dollars are spent on powerful biologics CD, but these powerful anti-inflammatory therapies have not significantly altered the incidence of fibrostenosing CD.^4^ Currently, there are no accepted medical anti-fibrotic therapies for CD, making the development of medical anti-fibrotic a research priority.

A significant barrier in the development of anti-fibrotic therapies is an incomplete understanding of the pathophysiology of IBD-related intestinal fibrosis. Our understanding is further limited by a lack of *in vivo* models of intestinal fibrosis that are reliable and appropriately mimic the human condition. Murine models have been a mainstay of *in vivo* modeling of intestinal inflammation. It is notable that CD-associated fibrosis is primarily limited to the terminal ileum, and most murine models of IBD involve colonic inflammation^5^. This represents a significant limitation to the study of intestinal fibrosis in CD. Currently there is only one described spontaneous model of ileal inflammation and fibrosis, and this strain (SAMP1/YitFc) has significant breeding limitations which hamper its use.^6^ This represents an opportunity to enhance our understanding of fibrostenosing CD through development of new *in vivo* model systems that closely mimic the complex pathophysiology at play in human CD and are also highly reliable, reproducible, and are amenable to both mechanistic and translational applications.

The TNF^ΔARE^ mouse is a well described murine model of spontaneous inflammation that develops Crohn’s-like ileitis and a number of extraintestinal manifestations that mimic CD.^7, 8^ Due to a deletion of AU-rich elements (ARE) in the 3’ untranslated region of the tumor necrosis factor (*Tnf)α* mRNA transcript, *Tnfα* mRNA has greater stability and thus overproduction systemically. This overproduction of *Tnfα* leads to highly penetrant ileal inflammation that is localized to the terminal ileum, as well as inflammatory arthritis.^7–10^ Typically, in heterozygous mice the changes of ileitis begin to occur at 6 to 8 weeks old.^7, 9, 10^ Homozygous mutations lead to early demise, either *in utero* or shortly after birth.

Given the highly biomimetic characteristics of the TNF^ΔARE+/−^ mouse to CD, we sought to determine if these animals represent an appropriate model of ileal fibrosis observed in a large number of human CD patients. Histologic and biochemical analysis of TNF^ΔARE+/−^ tissues identify an age-related increase in biomarkers of overt fibrosis, including fibrosis-associated gene expression, disorganized collagen organization and overt ileal strictures in older mice. These studies identify the TNF^ΔARE+/−^ mouse model as a relevant and tractable model of intestinal fibrosis with potential use in uncovering novel mechanisms of fibrostenotic disease in human CD.

## METHODS

All authors had access to the study data and have reviewed and approved the final manuscript.

### Animal handling

We utilized the C57BL/6-TNF^ΔARE^ previously described^7, 11^ and wild type littermates that were all bred in house. All animals were maintained and handled according to protocols approved by the Institutional Animal Care and Use Committee (IACUC).

### Ileum collection and processing

After euthanization per IACUC protocol, the abdominal cavity was carefully opened and the cecum, colon, and small bowel were identified. The small intestine, cecum, and proximal colon were then removed and a section spanning mid-small bowel to mid colon was then positioned next to a ruler and in most cases a photo was obtained. Removal of some peri-intestinal fat was performed for clarity of visualization and to be able to straighten the specimen. Typically, in TNF^ΔARE +/−^ animals the distal 3.0-4.0 cm of the ileum was grossly affected/diseased, and thus roughly 1.0-5.0 cm of ileum was utilized for analysis in our TNF^ΔARE +/−^ and TNF^ΔARE −/−^ control animals. Tissue was then used for RNA purification and ileal casting.

### Luminal casting

Once the ileum, cecum, and proximal colon were removed, they were processed, examined for gross stricturing and photographed. The ileum was then cut at the ileocecal junction and approximately 5-6 cm proximally with either a scalpel or sharp dissecting scissors. Any stool was gently flushed out using phosphate buffered saline (PBS) that was injected into the proximal end of the ileum tissue using a 3mL or 5mL syringe and needle that was between 22 and 26.5 gauge. Stool can also be gently expelled using blunt tweezers, with care taken not to damage the tissue.

An agarose-based compound was prepared by heating until it was liquid and flows freely. For these experiments we primarily used Epredia HistoGel™ (REF HG-4000-012), with some experiments using a 2% agarose composed of tap water mixed with Fisher BP160-500 molecular biology grade, low EEO/multipurpose agarose. In a separate area, cold PBS was placed into a holding tray (we use weigh boats) and placed on ice. Each tray was labeled with the sample for the animal. The cold PBS will serve to hold the casted ileum and preserve the tissue while the agarose hardens.

Where needed, a portion of terminal ileum was removed from the end for RNA or protein analysis using sharp dissecting scissors or scalpel. The distal end of the remaining ileum was securely sealed using either suture to tie off, or by clamping with a hemostat. If the hemostat was used, it was be carefully positioned and supported using blocks or by wedging between other tools so as to be upright keeping the ileum straight and without putting strain on the tissue. Two to three mL of agarose gel was then drawn into a 5mL syringe with 22-26.5 gauge needle (generally we select the largest bore needle we can that will fit into the lumen of the ileum, needles smaller than 26.5 gauge have difficulty passing the gel due to viscosity.) The proximal ileum was then gently grasped with either hemostats or flat tweezers, and the needle with agarose filled syringe attached was carefully inserted into the open lumen, taking extreme care not to puncture the wall of the ileum. The grasping hemostat was then gently tightened around the needle manually to create a seal, and the agarose gel was gently injected into the lumen until the ileum was completely filled, revealing any defects/strictures that were present.

The needle was gently removed while the grasping hemostat was then tightened to seal the distal end, and the entire piece was carefully placed into the cold PBS. It was manually held in the cold PBS for about 10 seconds, until the agarose was hard enough that the hemostat(s) can be removed and agarose does not leak out the end. The ileum was now casted, and it was left in the cold PBS for approximately 30-60 minutes to allow the gel to fully cure. Once cured and hardened, the ileum was removed, analyzed and photographed. Using very fine dissecting scissors, the ileum was gently cut lengthwise to flay open the ileum and reveal the cast within.

Great care must be taken to cut open the ileum and nudge the cast out of the flayed ileum without damaging the cast. The cast was then analyzed and photographed, and the ileal tissue was fixed in 10% neutral buffered formalin for further histological analysis.

### Measurements of casted ileum for determination of presence of stricture and ileal narrowing

All measurements were performed using (Fiji Is Just) ImageJ (Fiji). Measurement scales were set internally using a ruler that was imaged with each piece.

### Determination of luminal area

Luminal casting was performed as described elsewhere. The casts were then removed by cutting the ileum longitudinally and removing the cast within, attempted to keep cast as intact as possible. The area of each cast, and thus intraluminal area, was measured using imageJ. Area was then normalized to length of cast. All measurements were performed using (Fiji Is Just) ImageJ. Measurement scales were set internally for each image using a ruler that was imaged with each piece.

### Histology

Following casting and cast removal, the ileum was then flayed open and fixed in 10% neutral buffered formalin at room temperature or 4 degree for a minimum of 24 hours. Tissue was then processed via formalin fixation paraffin embedding (FFPE), and stained using hematoxylin and eosin (H&E), picrosirius red (PSR), and trichrome. Slides were imaged using an Olympus BX41 microscope with Spot Insight FireWire 2 megasample camera and processed using Spot 5.3 software. For polarized images of PSR stains, an Olympus polarizing filter was used.

### Picrosirius red (PSR) collagen calculation methods

For collagen quantification, ImageJ was used to analyze picrosirius red (PSR) stained ileal tissue. A representative field was captured on 20x. Using Fiji the area of collagen to length of ileum was measured. After conversion to 8-bit image, a threshold brightness was selected to most accurately capture all collagen luminescence whilst eliminating artifact as able, and this setting was used for analysis of all slides within each age group. The threshold used for age 6-10 weeks was 11, aged 14-15 weeks was 14, aged 18-24 was 10, and aged 24+ was 8. After adjusting the threshold, the area of wall and submucosa spanning from serosa to the crypt bases was selected and the area of collagen was recorded. The area was then normalized to length of ileum to determine the ‘area per length of ileum.’ For the comparison of fold change across age groups seen in figure 7, the average count/length of wild type was calculated as an average, and the fold change of each animal was calculated as (sample value – average of wild type)/average of wild type. This was analyzed as fold change to account for differences in thresholds used for analysis between groups, which was most likely made necessary due to variations in stain quality.

### Wall width measurements

For evaluation of wall layer thickness, H&E and/or PSR was utilized. The entire slide was examined and the thickest representative portion was centered into one field of view. Three measurements roughly equidistant where then obtained from this field of view using imageJ and average to generate a wall thickness value for the ileum of each animal, measured from the serosa to the crypt base

### Fibrosis scoring

For fibrosis scoring, each sample was graded using H&E and PSR in four domains on a scale from 0-3 or 0-1. These scores included:

*disorganization/disarray of collagen fibers*

- 0 – none, physiologically organized
- 1 – minimal disorganization, few small breaks
- 2 – moderately disorganized but structure still easy to see
- 3 – severely disorganized, no evidence of normal physiological pattern, collagen infiltrates muscular layers

*fibromuscular hyperplasia/ muscularization/muscular obliteration of the submucosa*

- 0 – none, normal muscle submucosa content
- 1 – mild increase in muscle content
- 2 – large amount of muscle in submucosa, affects the morphology but roughly normal structure
- 3 – muscle has obliterated the entire submucosal space

### Second Harmonic Generation (SHG) collagen calculation methods

Slides were de-paraffinzed and re-hydrated using xylene and progressive concentrations of ethyl alcohol. We then imaged using label-free second harmonic generation with a laser-scanning confocal microscope (LSM 780 spectral, Carl Zeiss, Thornwood, NY). Once images were obtained further analysis was performed using Fiji. Threshold was set to 50 to best eliminate artifact. We then measured the (red/collagen) area of the submucosa and wall, less the epithelium and normalized to ileum length (μm).

### Transcriptional analysis

Whole ileal was collected and snap-frozen in RNA*later*™ solution (Invitrogen ref AM7021) and later thawed and homogenized using a Bio-Gen Series PRO200 homogenizer. We then used bio Basic EZ-10 DNAaway mini-Prep RNA isolation kits (catalog # BS88136) were used to isolate RNA from the homogenate according to manufacturer’s instructions. Complementary DNA (cDNA) was then generated via reverse transcription using the iScript cDNA Synthesis kit (Bio-Rad) according to manufacturer’s instructions. We then used SYBR® Green (Applied Biosystems) for PCR analysis using the following primers:

*Β-actin* (mouse):

forward primer: 5’-AACCCTAAGGCCAACCGTGAA-3’

reverse primer: 5’-TCACGCACGATTTCCCTCTCA-3’

*Tnfα* (mouse):

forward primer: 5’-CCCTCACACTCAGATCATCTTCT-3’

reverse primer: 5’-GCTACGACGTGGGCTACAG-3’

*Il1β* (mouse):

forward primer: 5’-GGATGATGATGATAACCTGC-3’

reverse primer: 5’-CATGGAGAATATCACTTGTTGG-3’

*Tgfβ1*(mouse):

forward primer: 5’-GGATACCAACTATTGCTTCAG-3’

reverse primer: 5’-TGTCCAGGCTCCAAATATAG-3’

*Axl*(mouse):

forward primer: 5’-AGAGCTGAAGGAGAAACTAC-3’

reverse primer: 5’-TGGCAATTTTCATGGTCTTC-3’

QuantStudio^™^ machine and software were used to perform qRT-PCR and analyzed as fold change to control using ΔΔCT method and β-actin as the housekeeping gene^12^.

## Statistical analysis

Statistical analysis was performed using GraphPad Prism. Depending on the data, an unpaired t-test (Welch’s where appropriate), one-way ANOVA, receiver operating characteristic (ROC) curve analysis, or binomial testing was used. Results were deemed statistically significant when p < 0.05.

## RESULTS

### Structural fibrotic changes of intestinal fibrosis

The presence of intestinal stricturing and luminal narrowing are some of the most significant clinical sequelae of Crohn’s disease.^13, 14^ We thus sought to determine if clinically analogous findings developed in the terminal ileum of aged TNF^ΔARE +/−^ mice.

### Presence of strictures in the terminal ileum of aged (24+ week) TNF^ΔARE +/−^ mice

Determining the presence of a stricture in human IBD patients is a clinical challenge.^15^ Examination of surgical specimens remains the most concrete method of determining the presence of strictures, but the use of cross-sectional imaging has emerged as another useful tool.^16^ There is a great deal of heterogeneity in interpretation of what constitutes a stricture on cross-sectional imaging based on luminal diameter, full bowel diameter, up-stream dilation, and wall thickness.^16^ To determine presence of strictures, we evaluated the terminal ileum of wild type and TNF^ΔARE +/−^ mice *ex vivo* using two methods. This initial analysis was inclusive of n=6 wild type and n=7 TNF^ΔARE +/−^ mice at 24+ weeks of age.

We manually inspected the terminal ileum both before and after injecting with an agarose-based gel as described in the *Luminal casting* methods section. This method was needed to distend the intestines fully for accurate assessment, as distension was recognized as an important feature for accurate determination of stricture in the small intestine.^17^ We found that the terminal ileum of TNF^ΔARE +/−^ mice aged 24+ weeks contain gross stricture(s) with upstream dilation, ragged and uneven contours. Once the luminal cast was removed, these features are again demonstrated, indicating the lumen contains strictures with upstream dilation and overall decreased luminal volume. Wild type ileum are smooth and uniform with no stricture nor upstream dilation. (**Fig. 1 A and B**). We then counted the number of strictures observed on manual inspection of specimens, analogous to surgical determination of presence of stricture. No strictures were detected in wild type specimens compared to an average of 1.14 ± (0.90) strictures detected in aged TNF^ΔARE +/−^ specimens, (P = 0.015) (**Fig. 1D**).

**Figure 1.**
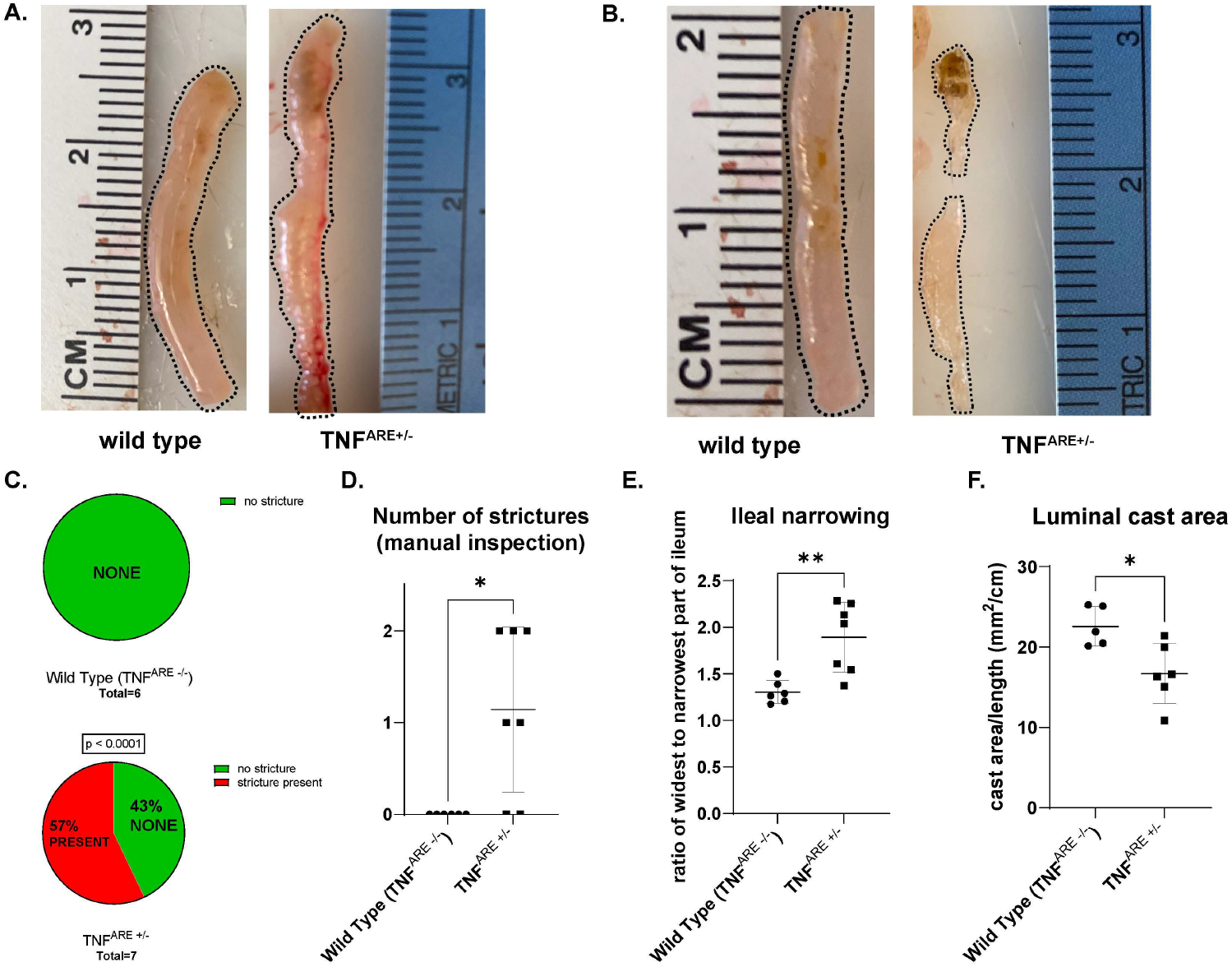
Aged (24+ week) TNF^ΔARE+/−^ mice develop clinically analogous structural fibrosis including stricture and luminal narrowing while age-matched littermate control wild type mice do not. These data are inclusive of 6 wild type and 7 TNF^ΔARE+/−^ mice. **A)** Terminal ileum that has been filled with agarose casts. **B)** Casts from 1A that have been removed from the ileum. **C)** Quantification of gross ileaI strictures in aged TNF^ΔARE+/−^ and wild type controls. **D)** Manual inspection of gross strictures in aged TNF^ΔARE+/−^ and wild type controls. E) Ratio of the width of the widest section to narrowest section of terminal ileum of WT controls and aged TNF^ΔARE+/−^ mice. F) Luminal cast area/length vvas measured. Data is expressed as mean ± SD. * P < 0.05, ** P < 0.01

We then sought to more objectively quantify the number of strictures present. All measurements were performed in terminal ileum that had undergone luminal casting as described in the *Luminal casting* methods section. Again, this method was needed to distend the intestines fully for accurate measurement of diameter, as distension was recognized as an important feature for accurate determination of stricture in the small intestine.^17^ To the authors knowledge, no such consensus on stricture determination exists for murine models. For this study, we extrapolated expert consensus from human imaging to create a definition of ileal stricture. An expert consensus on definition of naïve small bowel stricture in humans suggests that stricture be defined as luminal narrowing with bowel wall thickening and upstream dilation, with luminal diameter reduced by at least 50% relative to an adjacent bowel loop.^17^ Using Fiji, measurements were obtained to determine whether the terminal ileum contained any stricture meeting the rigorous and clinically analogous parameter of a narrowing with diameter ≤ 50% of an adjacent loop, and found that aged TNF^ΔARE +/−^ mice developed stricture in 57% of animals (P < 0.0001) compared to 0% wild type (**Fig. 1C**). To further characterize the ileal narrowing we compared the ratio of widest to narrowest measurement, and found that aged TNF^ΔARE +/−^ have significant ileal narrowing compared to wild type controls. The ratio of the width of the widest section to narrowest section of terminal ileum of wild type compared to aged TNF^ΔARE +/−^ mice was 1.3 ± (0.12) versus 1.9 ± (0.37) respectively (P = 0.005) (**Fig. 1E**).

### Presence of luminal narrowing in the terminal ileum of aged (24+ week) TNF^ΔARE +/−^ mice

We next sought to determine if luminal area, as a surrogate for luminal volume, was different in aged TNF^ΔARE +/−^ mice. Once the luminal casts were removed, the area per length of each cast was calculated using Fiji. These data show significantly reduced luminal area and thus luminal volume in aged TNF^ΔARE +/−^ mice compared to wild type controls. Luminal cast area quantified as area/length demonstrates wild type mice mean area of 22.57 ± (2.46) mm^2^/cm compared to 16.72 ± (3.72) mm^2^/cm in aged TNF^ΔARE +/−^ mice, (P = 0.015) (**Fig. 1F**).

These results demonstrate that at 24+ weeks of age, there are structural changes in the terminal ileum of TNF^ΔARE +/−^ mice such as stricture and luminal narrowing that are clinically analogous to the intestinal fibrosis seen in human CD patients.

### Histological features of intestinal fibrosis

Histologic evidence of intestinal fibrosis is difficult to characterize. Based on histologic features identified in these studies as important indicators of fibrosis in human Crohn’s disease,^18, 19^ we observed increased wall thickness (measured from serosa to mucosal layer/crypt base), increased collagen content/collagen expansion, disorganization/disarray of collagen fibers, fibromuscular hyperplasia and muscular obliteration of the submucosa as key features of small intestinal fibrosis.^18, 19^ We then created a fibrosis scoring system to capture features of disorganization/disarray of collagen fibers, fibromuscular hyperplasia and muscular obliteration of the submucosa, and performed objective measurements of wall thickness and amount of collagen content in the terminal ileum of TNF^ΔARE +/−^ mice.

### Wall width

We measured the wall width of the terminal ileum of wild type and TNF^ΔARE +/−^ mice after FFPE and staining with H&E and/or trichrome. The mean wall width of wild type and aged TNF^ΔARE +/−^ mice was 106.6 ± (26.61) μm versus 467.8 ± (171.30) μm respectively (P < 0.0001) (**Fig. 2A and B**). These data support the contention that the wall of the terminal ileum was significantly increased in aged TNF^ΔARE +/−^ mice.

**Figure 2.**
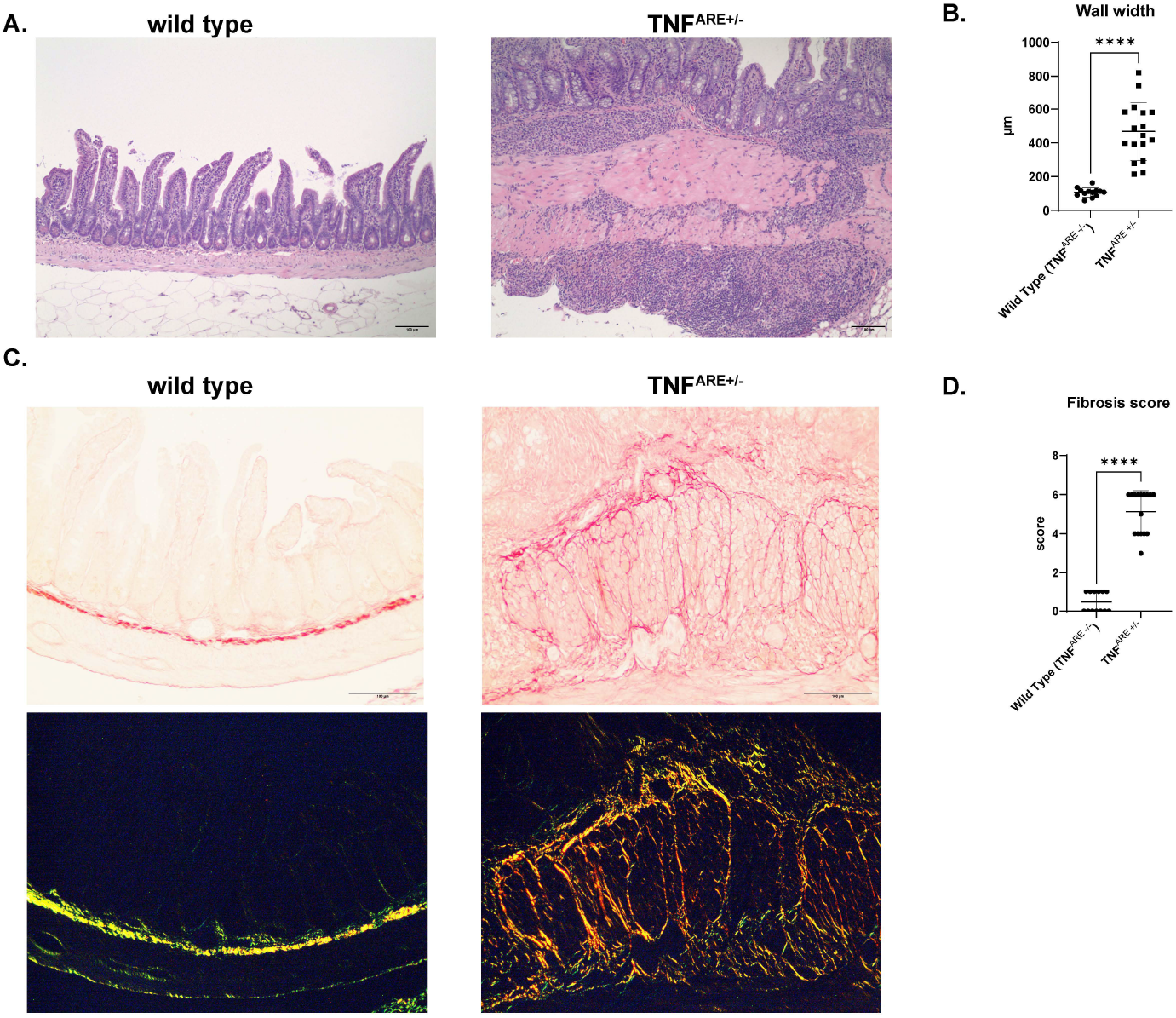
Aged (24+ week) TNF^ΔARE+/−^ mice develop histological fibrosis while age-matched littermate control wild type mice do not. **A)** Representative histology of terminal ileum of wild type and aged TNFMRE+t-using hematoxylin and eosin (H&E) staining, (n = 13 wild type and 16 TNF^ΔARE+/−^) **B)** Wall width measurements of wild type and aged TNF^ΔARE+/−^ mice. **C)** Representative histology of terminal ileum of wild type and aged TNF^ΔARE+/−^ using picrosirius red (PSR) staining. Top panel was under white light, lower panel was under polarized light. **D)** Fibrosis score of wild type and aged TNF^ΔARE+/−^ mice based on histological scoring. Data is expressed as mean ± SD. **** P < 0.0001

### Fibrosis score

To evaluate whether the TNF^ΔARE +/−^ mice develop histological changes of fibrosis consistent with human disease, we created a fibrosis scoring criteria using features published in a large systematic review and international consensus manuscripts.^18, 19^ For fibrosis scoring, each slide was graded using H&E and/or trichrome stain as well as PSR in two domains on a scale as described in the methods section.

At 24+ weeks of age, TNF^ΔARE +/−^ mice develop all of the above features of intestinal fibrosis in the terminal ileum. Representative images of H&E (**Fig. 2A**) and PSR under both white light and polarized light (**Fig. 2C**) demonstrate increased wall thickness, increased collagen content/collagen expansion, disorganization/disarray of collagen fibers, fibromuscular hyperplasia and muscular obliteration of the submucosa. The average fibrosis score of wild type to aged TNF^ΔARE +/−^ mice was 0.46 ± (0.52) to 5.13 ± (1.09) respectively (P < 0.0001) (**Fig. 2E**). These data support that substantial fibrotic changes occur histologically in aged TNF^ΔARE +/−^ mice after 24 weeks of age.

#### Collagen content in the terminal ileum of aged TNF^ΔARE +/−^ mice

While the above data show disorganization of collagen, we also objectively evaluated changes in collagen content. To do this, collagen content was quantified using second harmonic generation (SHG). Using SHG imaging, we quantified collagen amount in a sub-set of our wild type mice versus aged TNF^ΔARE+/−^. These data included 20 mice, 10 per group, and showed average collagen area (μm^2^) per length (μm) to be 24.28 ± (7.25) in the wild type and 67.05 ± (40.38) in the aged TNF^ΔARE +/−^ (P = 0.0085) (**Fig. 3**).

**Figure 3.**
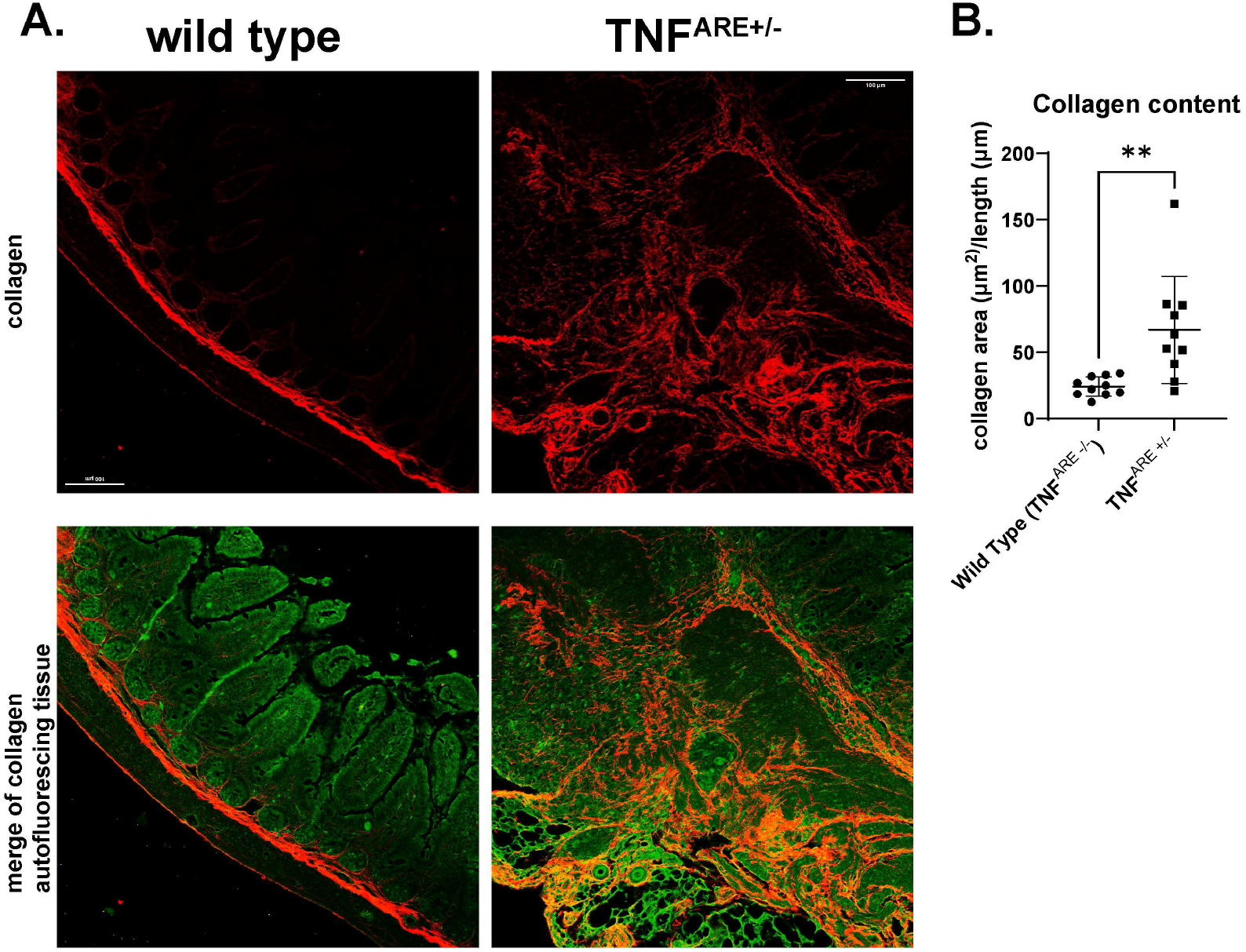
The increased collagen content deposition of aged TNF^ΔARE+/−^ mice in the terminal ileum wall is shown using second harmonic generation (SHG) on a subset of our 24+ week age matched littermate controls and aged TNF^ΔARE+/−^ mice. Representative images are shown. These data are inclusive of 10 age-matched littermate control wild type and 10 aged TNF^ΔARE+/−^ mice. **A)** Representative images of the terminal ileum of wild type and aged TNF^ΔARE+/−^ mice. **B)** Collagen content in the ileal wall quantified using SHG and reported as pixels of collagen (count) per length. Data is expressed as mean± SD. ** P < 0.01

These results demonstrate that at 24+ weeks of age TNF^ΔARE +/−^ mice have developed the key features of histological fibrosis in the terminal ileum that mimic classical features of intestinal fibrosis in human CD patients.

### Gene expression

#### Pro-inflammatory and pro-fibrotic gene expression in the terminal ileum of aged and young TNF^ΔARE +/−^ mice

After determining that structural and histological fibrosis develops in TNF^ΔARE +/−^ mice over 24 weeks old, we next sought to determine if select pro-inflammatory and pro-fibrotic gene expression was increased. RNA from whole tissue samples of terminal ileum was purified and analyzed with qPCR (wild type n=8, TNF^ΔARE +/−^ n=10) (**Fig. 4A**). Aged TNF^ΔARE +/−^ mice demonstrate a 29.37 ± (16.68)-fold increase compared to wild type in *Tnfα* mRNA expression (p = 0.0005), and a 37.89 ± (26.74)-fold increase compared to wild type in *Il1β* expression (P = 0.002). This analysis confirmed that *Tnfα* was appropriately elevated, and the increase in *Il1β* further demonstrates that the terminal ileum was in an inflamed state. Aged TNF^ΔARE +/−^ mice also demonstrate a 5.11 ± (2.39)-fold increase compared to wild type in *Tgfβ1* expression (P = 0.0004), and a 2.85 ± (0.79)-fold increase compared to wild type in *Axl* expression (P < 0.0001). *Tgfβ1* is well recognized as a pivotal regulator of fibrosis,^20–23^ and *Axl* has been shown to be a regulatory pathway in fibrosis in many organs including the small intestine.^24^ Thus, two pivotal pro-inflammatory genes and two pro-fibrotic genes showed increased expression in TNF^ΔARE +/−^ mice at an age of >24 weeks.

**Figure 4.**
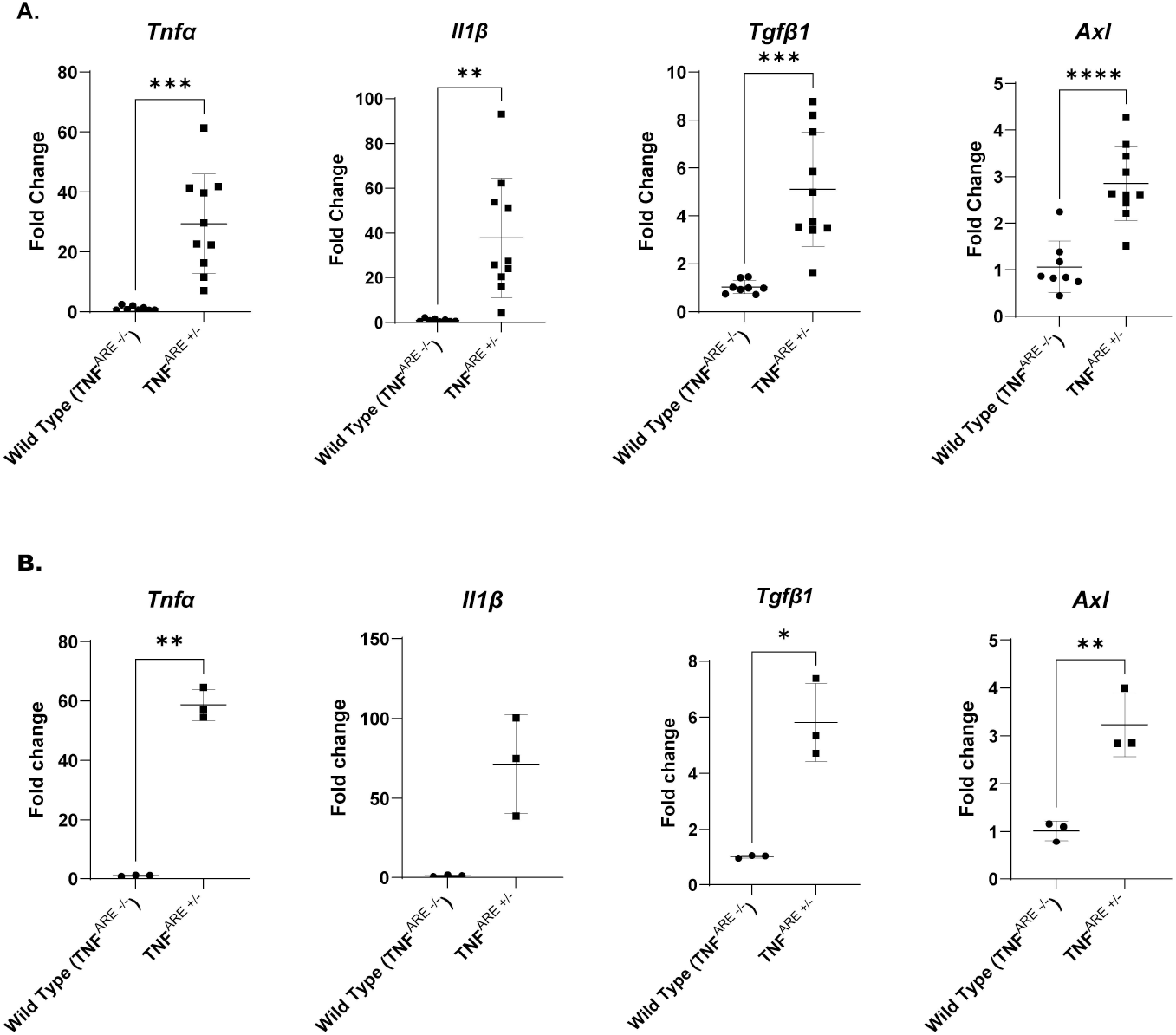
Aged TNF^ΔARE+/−^ mice express increases in several pro-inflammatory and pro-fibrotic genes in the terminal ileum, and these changes occur as early as 6 weeks. **A)** Gene expression in 24+ week old mice. These data are inclusive 9 age-matched wild type litter mate controls and 10 TNF^ΔARE+/−^. B) Gene expression in 6 week old mice. These data are inclusive of 6 mice, 3 age-matched wild type litter mate controls and 3 TNF^ΔARE+/−^. Data is expressed as mean ± SD. * P < 0.05, ** P < 0.01, *** P < 0.001, **** P < 0.0001

To determine if these genes represent appropriate biomarkers of ileal fibrosis, we utilized receiver operating characteristic (ROC) curves to analyze for association between fold-change of gene expression and fibrosis score as detailed above in 24+ week old TNF^ΔARE +/−^ mice (supplemental Figure 1). These analyses show a strong association between fibrosis score and increased gene expression of *Tnfα* (P = 0.0002), *Il1β* (P = 0.0005), and *Axl* (P = 0.002). No association was found with *Tgfβ1*, we hypothesize this was due to the complexity of Tgfβ1 signaling and heterogeneity of expression in our mice. Ultimately the ROC analysis supports use of these genes as markers for tissue fibrosis in 24+ week old TNF^ΔARE +/−^ mice.

To determine if these expression profiles occur at younger ages, we then analyzed mice at 6 weeks of age (wild type n=3, TNF^ΔARE +/−^ n=3). The increased expression of *Tnfα, Tgfβ1*, and *Axl* was present as early as 6 weeks, and, *Il1β* was increased substantially but narrowly missed statistical significance. (**Fig. 4B**). Six-week old TNF^ΔARE +/−^ mice demonstrate a 58.8 ± (5.23)-fold increase in *Tnfα* mRNA expression (P = 0.0027), and a 71.5 ± (30.88)-fold increase in *Il1β* expression (P = 0.06). Six-week old TNF^ΔARE +/−^ mice also demonstrate a 5.8 ± (1.39)-fold increase in *Tgfβ1* expression (P = 0.03), and a 3.2 ± (0.67)-fold increase in *Axl* expression (P = 0.0053).

Thus, it appears that the genetic milieu that ultimately results in fibrosis may be present as early as 6 weeks.

#### Profiling the progression to ileal fibrosis in TNF^ΔARE +/−^ mice

##### Age of emergence of histological fibrotic changes in terminal ileum of TNF^ΔARE +/−^ mice

Given the young age in which pro-fibrotic gene expression occurs, we next sought to characterize at what age structural and histologic fibrosis emerges in the ileum of TNF^ΔARE +/−^ mice. To determine at what age histological fibrosis emerges in the terminal ileum, we repeated the H&E, PSR, collagen quantification, wall width measurements, and fibrosis scoring performed on 24+ week old mice on three younger age cohorts. These cohorts are aged 7-10 weeks (wild type = 4, TNF^ΔARE +/−^ = 3), 14-15 weeks (wild type = 5, TNF^ΔARE +/−^ = 5), and 18-24 weeks (wild type = 3, TNF^ΔARE +/−^ = 3). Representative images of H&E and PSR staining under white light and polarized light show that at 7-10 weeks, TNF^ΔARE +/−^ terminal ileum was nearly indistinguishable from wild type, but pro-inflammatory and pro-fibrotic are seen at 14-15 weeks (**Fig. 5A and B**). These are visually less pronounced than that was seen in animals aged over 24 weeks.

**Figure 5.**
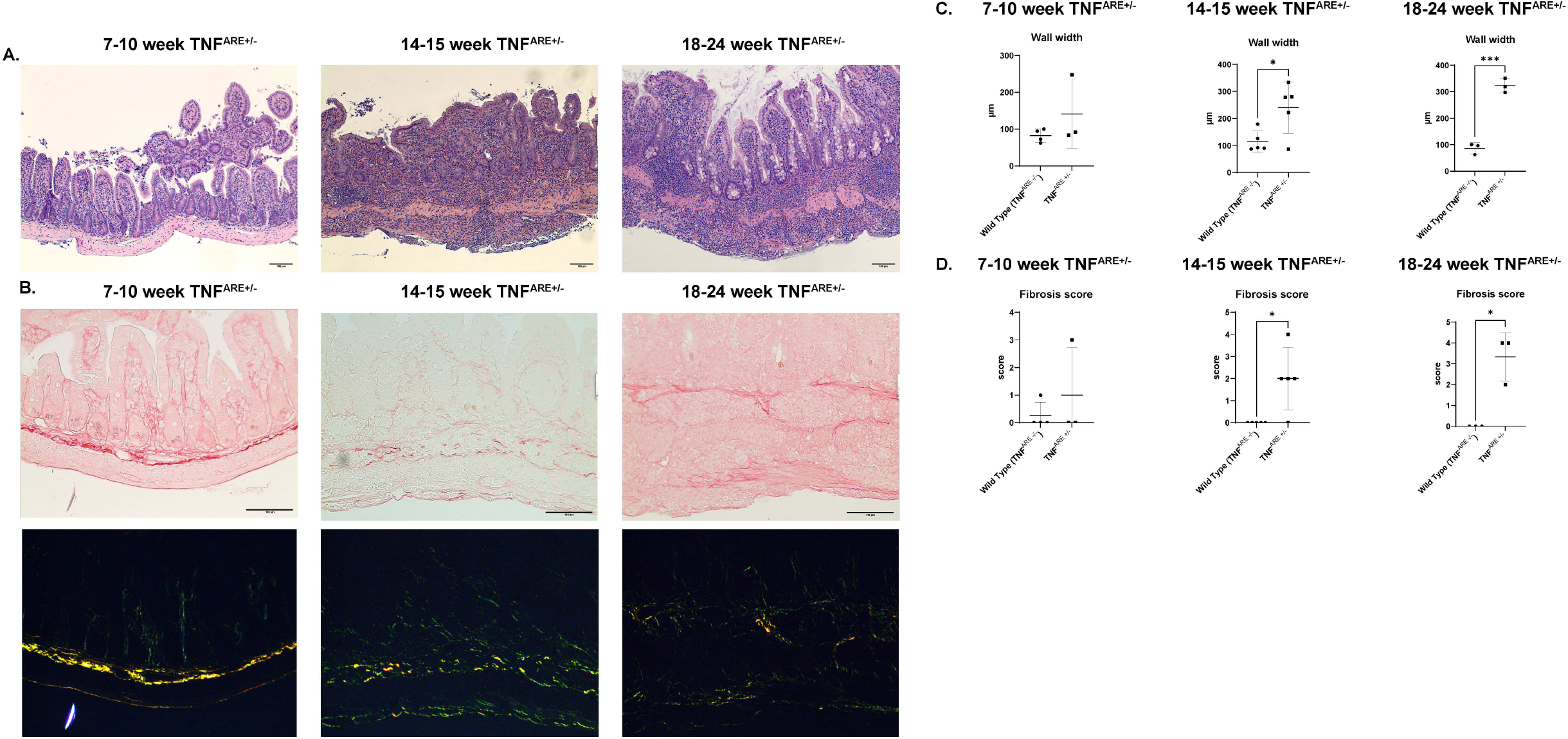
Age-related histological changes of intestinal fibrosis. **A)** Representative histology of terminal ileum of wild type and aged TNF^ΔARE+/−^ using hematoxylin and eosin (H&E) staining. **B)** Representative histology of terminal ileum of wild type and TNF^ΔARE+/−^ of varying ages using picrosirius red (PSR) staining. Top panel was under white light, lower panel was under polarized light. **C)** Wall width measurements of wild type and TNF^ΔARE+/−^ mice aged 7-10 weeks, and aged 18-24 weeks. **D)** Fibrosis score based on histological scoring of wild type and TNF^ΔARE+/−^ mice aged 7-10 weeks, aged 14-15 weeks, and aged 18-24. Data is expressed as mean± SD.* P <0.05, P < 0.001

The wall width was not increased in TNF^ΔARE +/−^ mice aged 7-10 weeks, but was significantly thicker in animals aged 14-15 weeks and 18-24 weeks (**Fig. 5C**). Wall width measurements of wild type compared to TNF^ΔARE +/−^ mice at 7-10 weeks was 82.6 ± (17.9) μm and 141.3 ± (92.3) μm respectively (P = 0.39), at 14-15 weeks was mean width 114.7 ± (39.2) μm and 240.0 ± (94.3) μm respectively (P = 0.03), and at 18-24 weeks was mean width 86.45 ± (20.9) μm and

322.9 ± (26.5) μm respectively (P = 0.0003). Thus, wall width begins to increase by the 14-15 week. This was likely due to a combination of developing fibrosis but also strongly influenced by immune cell infiltration, which is a characteristic of the TNF^ΔARE +/−^ mice by this age.

The fibrosis score based on the histological scoring described in methods of wild type compared to TNF^ΔARE +/−^ mice was unchanged in the youngest cohort, but was significantly increased compared to wild type in the 14-15 week and 18-24-week cohorts (**Fig. 5E**). Wild type to TNF^ΔARE +/−^ scores aged 7-10 weeks were [mean scores 0.25 ± (0.50) and 1 ± (1.73) respectively (P = 0.44)], aged 14-15 weeks [mean score 0 ± (0.00) and 2 ± (1.41) respectively (P = 0.034)], and aged 18-24 weeks [mean score 0 ± (0.00) and 3.33 ± (1.12) respectively (P = 0.038)].

These data demonstrate that at age 7-10 weeks, TNF^ΔARE +/−^ mice do not exhibit any histological evidence of fibrosis. Beginning in around 14-15 weeks, wall thickening and fibrotic changes begin to occur and this continues through the 14=15 week and 18-24-week age range. Collagen content does not increase in the pre-24-week animals compared to wild type.

##### Age of emergence of structural fibrotic changes in the terminal ileum of TNF^ΔARE +/−^ mice

To determine at what age clinically analogous strictures emerge we repeated the ileal casting and measurements performed on 24+ week old mice on two younger age cohorts. These cohorts are aged 7-10 weeks (wild type = 4, TNF^ΔARE +/−^ = 3) and aged 16-20 weeks (wild type = 3, TNF^ΔARE +/−^ = 4). Grossly, the terminal ileum did not demonstrate any strictures by observation or measurement in the 7-10-week aged cohort (**Fig. 6A, B and D**). Further, 7-10-week old TNF^ΔARE +/−^ mice do not develop ileal narrowing (P = 0.86), nor smaller luminal area volumes (p = 0.13) (**Fig. 6C and E**). At the age of 16-20 weeks, TNF^ΔARE +/−^ mice do have some nearly significant observed evidence of gross ileal strictures (mean of 0.75 ± (0.50) strictures in TNF^ΔARE +/−^ mice, none in wild type, p = 0.06) (**Fig. 6F, G and I**), but no ileal narrowing (P = 0.70) (**Figure 6H**), nor smaller luminal area volumes (**Figure 6J**) (P = 0.13).

**Figure 6.**
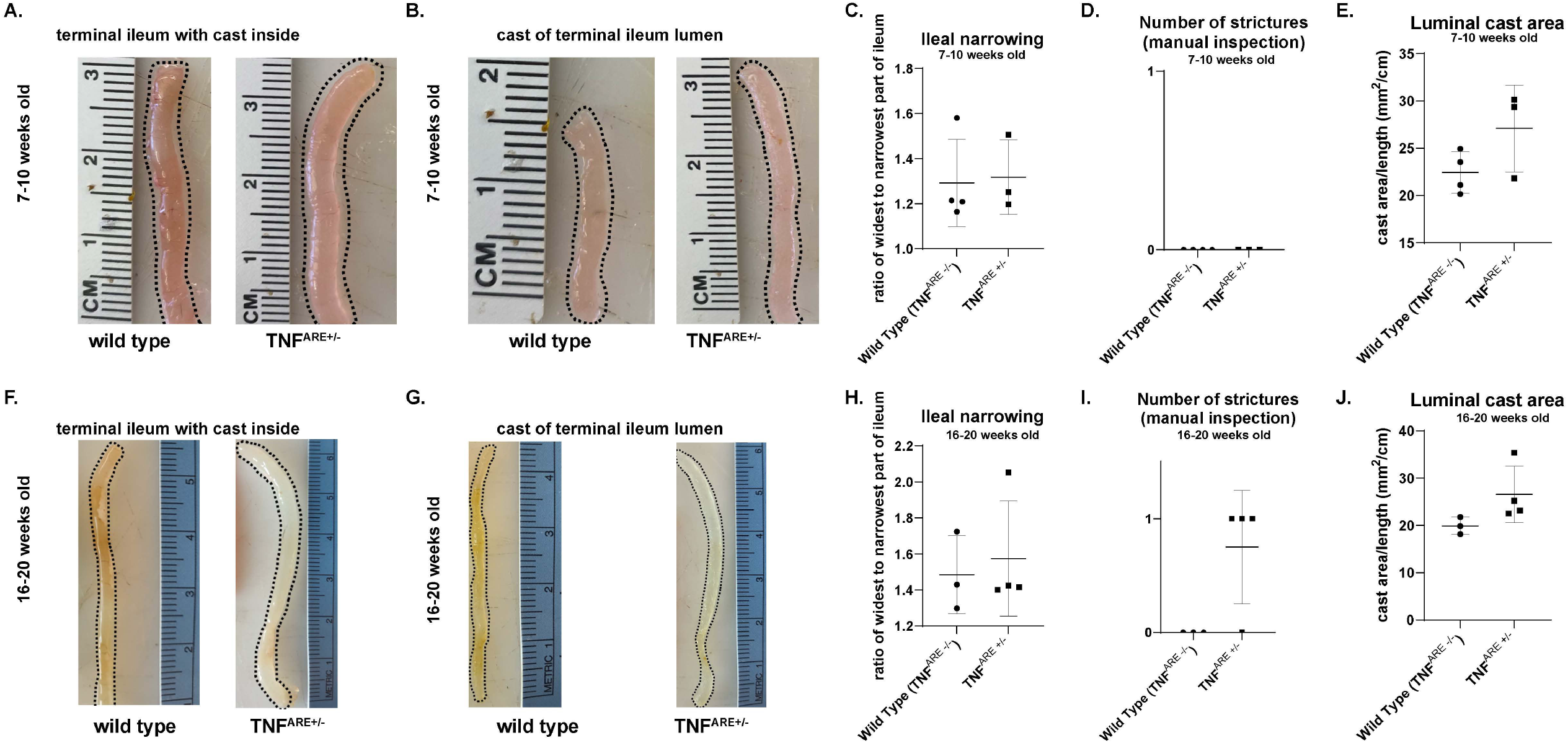
TNF^ΔARE+/−^ mice do not develop robust clinically analogous structural fibrosis until after 20 vveeks of age. These cohorts are aged 7-10 weeks **(A-E)** (n for wild type= 4 and TNF^ΔARE+/−^ = 3) and aged 16-20 weeks **(F-J)** (n for wild type= 3 and TNF^ΔARE+/−^ = 4). **A and F)** Terminal ileum that has been filled with agarose casts at ages 7-10 weeks **(A)** and 16-20 weeks **(F)**. Once removed, the casts of the same mice is shown in **Band G)** at ages 7-10 vveeks **(B)** and 16-20 weeks **(G). C-E)** Quantification of gross ileal strictures, ileal narrowing, and smaller luminal area volumes in wild type and 7-10-week old TNF^ΔARE+/−^ mice H-J) Quantification of gross ileal strictures, ileal narrowing, luminal area volumes in wild type and 16-20-week old TNF^ΔARE+/−^-mice.

Thus, there was evidence that some degree of ileal stricturing was beginning to develop at 16-20 weeks of age, but there was no significant change in ileal narrowing, and no change in luminal cast area between wild type and TNF^ΔARE +/−^ mice. These results suggest that TNF^ΔARE +/−^ mice do not begin to develop clinically analogous stricturing until around 16-20 weeks of age, but more robust structural fibrosis occurs after 20 weeks.

#### Comparison of clinically analogous structural changes of ileal fibrosis across age groups

Having discovered that TNF^ΔARE +/−^ over 24 weeks of age exhibit several aspects of structural ileal fibrosis not seen in younger TNF^ΔARE +/−^ mice, we sought to establish a time-course for the development of ileal fibrosis in TNF^ΔARE +/−^ mice. For each of these, <u>we are showing the same data already presented above that have been re-organized for direct comparison.</u> For each of the following endpoints examined, the wild-type cohorts of each age group were compared to each other and showed no significant difference (supplemental figure 2). We then pooled wild type data into one group regardless of age and compared to TNF^ΔARE +/−^ mice of each age cohort. Structural parameters of observed strictures, ileal narrowing, and luminal cast area were compared.

TNF^ΔARE +/−^ mice do not develop significant gross strictures until 24+ weeks (**Fig. 7A**). The mean number of strictures in wild type, 7-10-week old TNF^ΔARE +/−^ mice, 16-20-week old TNF^ΔARE +/−^ mice, and 24 + week old TNF^ΔARE +/−^ mice was 0 ± (0.00), 0 ± (0.00), 0.75 ± (0.50, and 1.143 ± (0.90) respectively (P = 0.0003). There was a significant increase in number of strictures in 24+ week old TNF^ΔARE +/−^ mice compared to wild type (P = 0.0003) and compared to the youngest cohort of old TNF^ΔARE +/−^ mice (P = 0.01). No significant differences were seen when comparing wildtype to 7-10-week old or 16-20-week old TNF^ΔARE +/−^ mice, when comparing 7-10-week old to 16-20-week old TNF^ΔARE +/−^ mice, nor when comparing 16-20-week old to 24+ week old TNF^ΔARE +/−^ mice.

**Figure 7.**
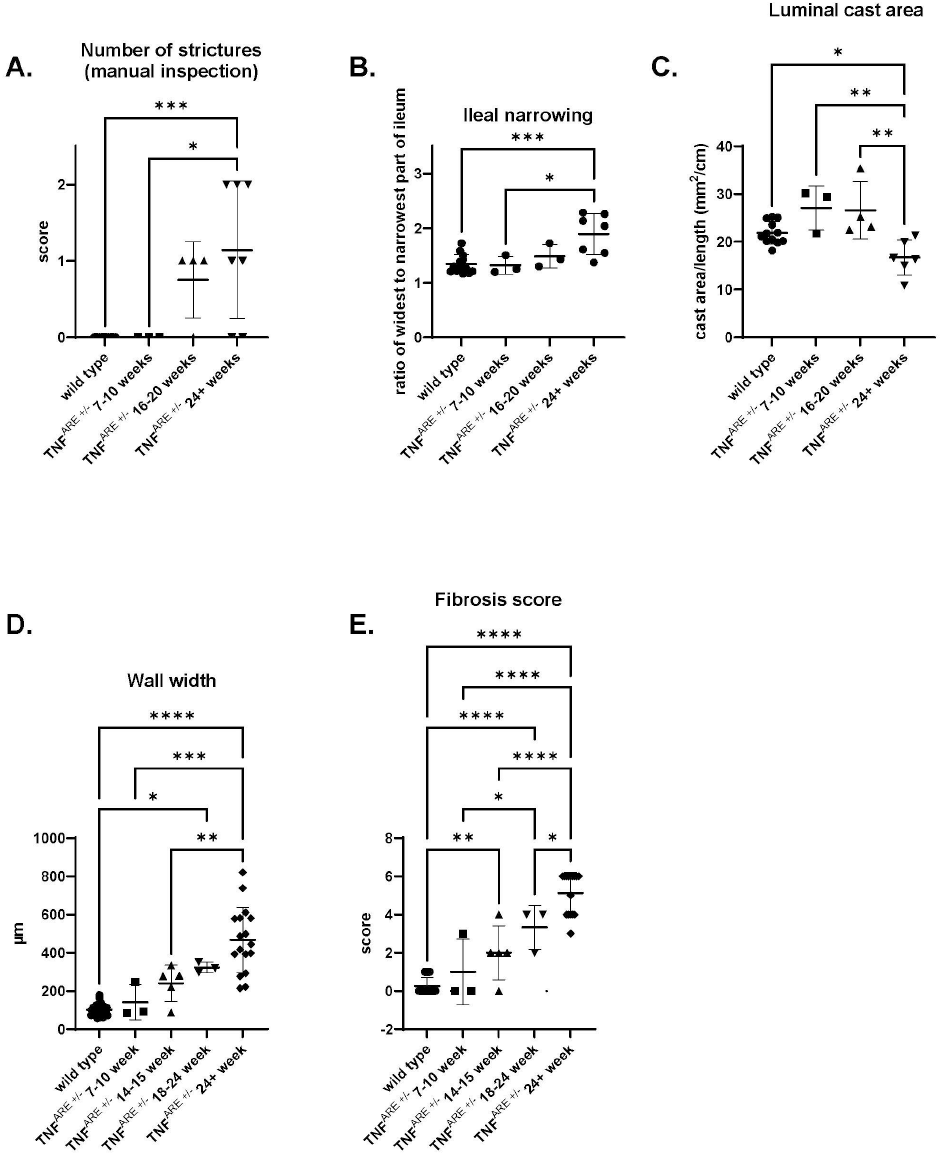
TNF^ΔARE+/−^ mice develop structural and histological changes of ileaI fibrosis over time. These data <u>are the same data already presented in other figures that have been re-organized for direct comparison.</u> All wild type across age groups has been pooled. These demonstrate the progression of number of ileal strictures **(A)**, ileal narrowing **(B)**, luminal cast area **(C)**, wall width **(D)**, and fibrosis score **(E)** in wild type and TNF^ΔARE+/−^ at 7-10, 16-20, and 24+ weeks. * p < 0.05, ** p < 0.01, *** p < 0.001, **** p < 0.0001

TNF^ΔARE +/−^ mice do not develop significant ileal narrowing until 24+ weeks (**Fig. 7B**). The mean ratio of widest to narrowest section of terminal ileum in 7-10-week old TNF^ΔARE +/−^ mice, 16-20-week old TNF^ΔARE +/−^ mice, and 24 + week old TNF^ΔARE +/−^ mice was 1.342 ± (0.17), 1.319 ± (0.17), 1.484 ± (0.22), and 1.892 ± (0.37) respectively (P = 0.0008). There was a significant increase in the ratio of widest to narrowest section of terminal ileum in 24+ week old TNF^ΔARE +/−^ mice compared to wild type (P = 0.0005) and compared to the youngest cohort of old TNF^ΔARE +/−^ mice (P = 0.01). No significant differences were seen when comparing wildtype to 7-10-week old or 16-20-week old TNF^ΔARE +/−^ mice, when comparing 7-10-week old to 16-20-week old TNF^ΔARE +/−^ mice, nor when comparing 16-20-week old to 24+ week old TNF^ΔARE +/−^ mice.

TNF^ΔARE +/−^ mice do not develop significant reductions in luminal cast area until 24+ weeks (**Fig. 7C**). The mean luminal cast area of terminal ileum in wild type, 7-10-week old TNF^ΔARE +/−^ mice, 16-20-week old TNF^ΔARE +/−^ mice, and 24 + week old TNF^ΔARE +/−^ mice was 21.87 ± (2.34) mm^2^/cm, 27.11 mm^2^/cm ± (4.61), 26.57 ± (5.99) mm^2^/cm, and 16.72 ± (3.73) mm^2^/cm respectively (P = 0.0008). There was a significant decrease in cast area (a surrogate for luminal volume) in 24+ week old TNF^ΔARE +/−^ mice compared to wild type (P = 0.05), compared to the youngest cohort of old TNF^ΔARE +/−^ mice (P = 0.003), and compared to the 16-20-week age group (P = 0.002). No significant differences were seen when comparing wildtype to 7-10-week old or 16-20-week old TNF^ΔARE +/−^ mice, nor when comparing 7-10-week old to 16-20-week old TNF^ΔARE +/−^ mice.

These data indicate that structural parameters of observed strictures, ileal narrowing, and luminal cast are do not significantly change until TNF^ΔARE +/−^ mice reach at least 24 weeks of age.

#### Comparison of histological changes of ileal fibrosis across age groups

Having discovered that TNF^ΔARE +/−^ over 24 weeks of age exhibit several histological aspects of ileal fibrosis that may not emerge until later in life, we sought to establish a time-course for the development of histological features of ileal fibrosis in TNF^ΔARE +/−^ mice. For each of these endpoints <u>we are showing the same data already presented above that have been re-organized for direct comparison</u>. For each of the following endpoints examined, the wild-type cohorts of each age group were compared and showed no significant difference (supplemental figure 1). We then pooled wild type data into one group regardless of age and compared to TNF^ΔARE +/−^ mice of each age cohort. Histological parameters of H&E and PSR staining, wall width, and fibrosis score were compared.

When comparing across all age groups, the ileal wall width of TNF^ΔARE +/−^ mice was significantly thicker at 18-24 weeks. The mean wall width of the terminal ileum of wild type, 7-10-week old TNF^ΔARE +/−^ mice, 14-15-week old TNF^ΔARE +/−^ mice, 18-24eek old TNF^ΔARE +/−^ mice, and 24 + week old TNF^ΔARE +/−^ mice was 101.9 ± (28.71) μm, 141.3 ± (92.31) μm, 240.0 ± (94.34) μm, 322.9 ± (26.50) μm, and 467.8 ± (171.3) μm respectively (P < 0.0001). There was a significant increase in wall width of the terminal ileum in 24+ week old TNF^ΔARE +/−^ mice compared to wild type (P < 0.0001), (7-10 weeks P < 0.0001), and (14-15 weeks P < 0.0001), but not the 18-24-week cohort). The 18-24-week age group had significantly increased score compared to wild type (P = 0.01). No other significant changes were seen. When combined with the data directly comparing wall width to wild type in the 14-15-week age group, these data demonstrate a progression of terminal ileum wall thickening that was not present in younger animals but begins in TNF^ΔARE +/−^ mice aged 14-24 weeks.

TNF^ΔARE +/−^ mice begin to develop observable histological evidence of fibrosis emerging at 14-15 weeks (**Fig. 7E)**. The mean fibrosis scores of wild type, 7-10-week old TNF^ΔARE +/−^ mice, 14-15-week old TNF^ΔARE +/−^ mice, 18-24-week old TNF^ΔARE +/−^ mice, and 24 + week old TNF^ΔARE +/−^ mice were 0.25 ± (0.44), 1 ± (1.73), 2 ± (1.41), 3.33 ± (1.16), and 5.125 ± (1.09) respectively (P < 0.0001). There was a significant increase in fibrosis score of the terminal ileum in 24+ week old TNF^ΔARE +/−^ mice compared to all other groups (wild type P < 0.0001, 7-10 weeks P < 0.0001, 14-15 weeks P < 0.0001, and 18-24 weeks P = 0.03). The 18-24-week age group had significantly increased score compared to wild type (P < 0.0001) and the 7-10-week age group (P = 0.03), but no difference compared to the 14-15-week cohort. The 14-15-week cohort was also significantly increased compared to the wild type cohort (P = 0.003) but not the 7-10-week cohort. No significant differences were seen when comparing wildtype to 7-10-week old cohort. These data demonstrate a progression of histological fibrosis development that is not present in younger animals but begins in TNF^ΔARE +/−^ mice aged 14-15 weeks.

These data indicate that histological parameters of ileal wall width and collagen disorganization with muscular hyperplasia (fibrosis score) do not significantly change until TNF^ΔARE +/−^ mice reach 14-15 weeks but are most pronounce after at least 24 weeks of age.

## DISCUSSION

There is significant interest in the development of anti-fibrotic therapies for diseases such as CD.^17, 25^ The field is currently limited by a lack of understanding of the pathophysiology of IBD-related intestinal fibrosis. A significant void is the existence of appropriate *in vivo* models that mimic human disease. Current models of intestinal fibrosis include the use of spontaneous mutations, targeted gene deletions, chemically-induced fibrosis and infectious inflammatory models.^6^ While each model provides valuable insight into the fibrotic response, each of the currently utilized models is also fraught with inconsistency and sporadic appearance of fibrotic lesions.^6^ The goal of the present study was to determine whether TNF^ΔARE +/−^ mice may provide insight into the development of structuring fibrosis, and as such, serve as a model to define some of these principles.

The TNF^ΔARE^ model was originally developed by targeted deletion of 69 base pairs of AU-rich elements in the 3’ UTR region of the TNF gene^7^. This deletion results in increased TNF mRNA stability, increased TNF protein synthesis and elevated systemic TNF levels. The disease is mediated by leukocytes that intensely infiltrate the intestinal lamina propria. T cells isolated from the mesenteric lymph nodes of TNF^ΔARE^ mice, but not those from C57BL6/J mice, produce high levels of TNF-α during early disease (3-4 weeks-of-age) and a mixed Th1:Th2 cytokine profile during the later stages of chronic ileitis. TNF^∆ARE^ T cells can adoptively transfer disease to naïve, non-predisposed RAG^−/−^ recipients. The adoptively transferred disease is different from that transferred by CD45Rb^high^ cells, as TNF^ΔARE+/−^ T cells predominantly induce ileitis, demonstrating an inherent capacity to recirculate preferentially to the small intestine.^8^ Lastly, TNF^ΔARE^ mice develop intestinal lymphangiectasia, lymphangiogenesis, and tertiary lymph node organs in the mesentery.^26^ These features bear great similarity to the lymphatic changes seen in human CD.^26–29^ Important in this regard, the TNF^ΔARE^ mouse recapitulates many features of human CD, including: 1) the spontaneous nature and chronic course of the disease, 2) its pathogenetic dependence on T cells and proinflammatory cytokines, 3) its specific small intestinal (ileal) localization, and 4) abnormal lymphatics with development of mesenteric tertiary lymphoid organs.

Our analysis of TNF^ΔARE +/−^ mice revealed development ileal fibrosis similar to what is seen in CD patients. As observed in human patients, there was a time-dependent component, and chronic inflammation preceded the development of fibrosis in these mice. The progression of pro-inflammatory gene induction occurred at a time point prior to 6 weeks, with the first signs of histological fibrosis appearing around 14-15 weeks, with profound histological and structural fibrosis occurring at >24 weeks of age. These analyses have several strengths. Our investigations and outcome measures were specifically designed based on the fibrosis that is seen in human patients. While histological features of intestinal fibrosis can be difficult to determine, we incorporated multiple aspects of intestinal fibrosis including wall thickness, disorganization of collagen, fibromuscular hypertrophy and obliteration, and increased collagen content. TNF^ΔARE +/−^ mice develop clear fibrotic changes in all of aspects of analysis.

Clinically analogous outcomes of intestinal stricture, decreased luminal area, and luminal narrowing were specifically evaluated. This was enabled by the use of an agarose-based casting technique that we developed to allow for characterization of these outcomes. This lends relevance to this model as a biomimetic of human disease. This casting technique was also transferrable to the study disease in any part of the gastrointestinal tract, and may also be adaptable to the study of other organs. The observation of a progression of changes at distinct ages, starting with gene expression and progressing through histological changes and lastly structural changes, lends itself to tailored study of interventions ranging from preventative therapy to agents aimed at reversal of fibrosis.

Although the TNF^ΔARE +/−^ model has many strengths, there are also some limitations. While in the heterozygote state, these mice are consistently viable and more robust than for some genetic models, there are logistical limitations due to the need to age the mice to see full intestinal fibrosis. Moreover, these animals did not develop overt intestinal stricture, fistulization and life-threatening obstruction often observed in CD patients.^3, 30^ It is possible that if allowed to age long enough these could develop, however our 24+ week cohort did contain multiple animals between 30-35 weeks of age and these changes were not seen. TNF^ΔARE +/−^ mice meet humane endpoints for termination prior to seeing these changes. It will be interesting to determine if this phenotype could be enhanced or progressed. It is possible, for example, that the combinatorial use of chemical fibrosis inducers (e.g. bleomycin)^31–33^ in the TNF^ΔARE+/−^ could provide earlier and more aggressive fibrotic endpoints. More work will be needed to uncover the complex mechanisms of fibrosis in this model. While the genetic modification is mono-genetic, the subsequent increases in other pro-inflammatory as well as pro-fibrotic cytokines suggest that multiple pathways become involved, and this is also corroborated by the known complexity and incomplete understanding of TNF signaling.^34, 35^

Taken together, this model has several clinically relevant, measurable, and targetable outcomes that can be studied. This model and the outlined techniques and outcome measures are conducive to the study of potential medical therapies as well as mechanistic studies in intestinal fibrosis. Thus, we posit the TNF^ΔARE +/−^ mice as a tractable and viable model for the study of intestinal fibrosis such as that seen in CD.

## Abbreviations

CD: (Crohn’s disease)
IBD: (inflammatory bowel disease)
UC: (ulcerative colitis)

---

**Figure S1.**
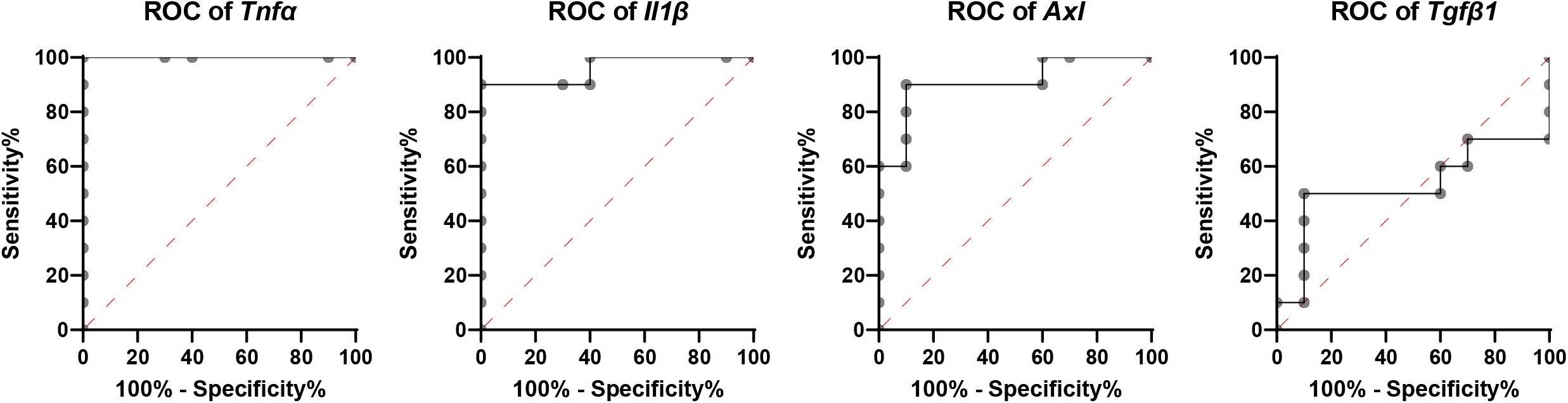
ROC curves showing gene expression correlation to fibrosis score for each of the four genes we analyzed.

**Figure S2.**
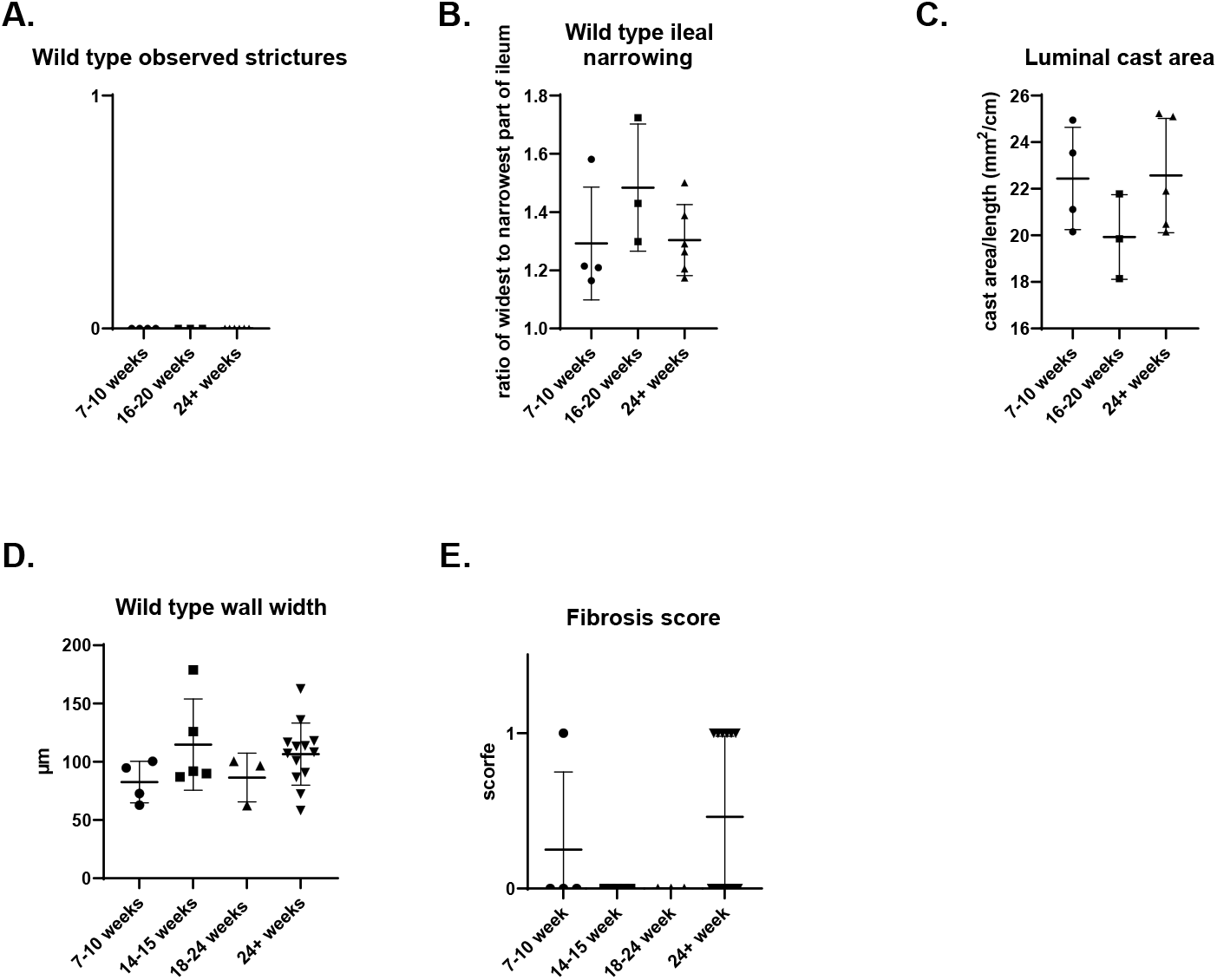
All wild type data from each age group for observed strictures (**A**), ileal narrowing (**B**), luminal cast area (**C**), ileal wall width (**D**), and fibrosis score (**E**). Wild types across all ages studied showed no difference in any of these parameters.

